# Structural diversity and clustering of bacterial flagellar outer domains

**DOI:** 10.1101/2024.03.18.585621

**Authors:** Jessie Lynda Fields, Hua Zhang, Nathan F. Bellis, Holly A. Petersen, Sajal K. Halder, Shane T. Rich-New, Hui Wu, Fengbin Wang

**Affiliations:** Department of Biochemistry and Molecular Genetics University of Alabama at Birmingham, Birmingham, AL 35233, USA; Department of Oral Rehabilitation & Biosciences Oregon Health & Science University, Portland, OR 97239, USA; Gregory Fleming James Cystic Fibrosis Research Center University of Alabama at Birmingham, Birmingham, AL 35233, USA

**Keywords:** bacterial flagella, flagellar filament, outer domains, Ig-fold, cryo-EM

## Abstract

Supercoiled flagellar filaments function as mechanical propellers within the bacterial flagellum complex, playing a crucial role in motility. Flagellin, the building block of the filament, features a conserved inner D0/D1 core domain across different bacterial species. In contrast, approximately half of the flagellins possess additional, highly divergent outer domain(s), suggesting varied functional potential. In this study, we elucidate atomic structures of flagellar filaments from three distinct bacterial species: *Cupriavidus gilardii*, *Stenotrophomonas maltophilia*, and *Geovibrio thiophilus*. Our findings reveal that the flagella from the facultative anaerobic *G. thiophilus* possesses a significantly more negatively charged surface, potentially enabling adhesion to positively charged minerals. Furthermore, we analyzed all AlphaFold predicted structures for annotated bacterial flagellins, categorizing the flagellin outer domains into 682 structural clusters. This classification provides insights into the prevalence and experimental verification of these outer domains. Remarkably, two of the flagellar structures reported herein belong to a previously unexplored cluster, indicating new opportunities on the study of the functional diversity of flagellar outer domains. Our findings underscore the complexity of bacterial flagellins and open up possibilities for future studies into their varied roles beyond motility.

## Introduction

More than 80% of all known bacterial species are motile at some stage of their life cycle^1^. These bacteria navigate toward nutrients or away from unfavorable environments through chemotaxis^2–4^, using movements such as swimming, swarming, and twitching, among others^5^. The flagellum, an extracellular machinery, facilitates swimming and swarming movements and comprises three main components: the basal body complex, functioning as the motor; the connecting rod and hook, acting as a universal joint; and the flagellar filaments, serving as a mechanical propeller^6,7^. Canonical flagellar filaments are long, supercoiled structures composed of around 20,000 flagellins. As the filament rotates, it generates thrust, acting similarly to an Archimedean screw. To date, all known bacterial flagellins share a conserved D0/D1 domain architecture, which is uniformly arranged across various bacterial species. These domains exhibit a helical rise of approximately 5 Å and a twist of 65.4 degrees, aligning every 11 flagellins near vertically. This configuration presents distinct 11-start protofilaments on the filament’s surface and its inner core. Recent research has revealed the molecular basis underlying flagellar supercoiling, identifying 11 different flagellin D0/D1 states that result in 11 unique protofilament conformations. Interestingly, similar supercoiling mechanisms have also been documented in archaeal flagella, despite the structural components of archaeal and bacterial flagella not sharing homology^8^.

For bacteria that colonize or infect other organisms, motility plays a crucial role in interactions between a bacterium and its host^5^. Beyond enabling movement, flagella are thought to possess several other functions, including adhesion to surfaces^9,10^, colonization^11^, biofilm formation^12^, and potentially modulating host immune response^13^, among others. Many of these properties are attributed to the flagellar outer domains, which are part of the central region of the flagellin, named D2, D3, D4, and so forth. In several bacterial species, these outer domains are not essential for motility^14,15^. In fact, about half of the bacterial flagellins annotated in the UniProt database contain only the D0/D1 domain, lacking outer domains.

Several outer domain structures from a diverse range of species^10,16–26^ that include soil-borne bacteria such as *Sinorhizobium meliloti*, opportunistic human pathobionts such as *Salmonella enterica* and *Pseudomonas aeruginosa*, as well as life-threatening primary human pathogens like *Burkholderia pseudomallei*, a high-priority biological agent responsible for melioidosis^27^, have been reported in the previous studies. These structures encompass flagellin structures solved by X-ray crystallography and filamentous structures determined by cryo-electron microscopy (cryo-EM). Remarkably, the fold and architecture of the outer domains are highly variable. The lack of clear homology among the outer domains of flagellin across different species has historically made it challenging in the structural analysis of these domains. Recent advances in protein structure prediction methods, notably AlphaFold^28^, have shown great promise in accurately predicting protein structures at the fold level, even when no analogous structure exists. With most UniProt sequences now predicted and available in the AlphaFold database^29^, it is feasible to undertake large-scale structural analyses of flagellin outer domains.

In this study, we report the near-atomic resolution cryo-EM structures of supercoiled bacterial flagellar filaments from three diverse bacterial species. The first is from *Cupriavidus gilardii*, a gram-negative, aerobic, opportunistic pathogen that has been increasingly associated with human infection and holds potential for bioremediation^30,31^. The second is from *Stenotrophomonas maltophilia*, a gram-negative, aerobic bacterium known for its multidrug resistance and its ability to infect the lungs of individuals with cystic fibrosis^32,33^. The third is from *Geovibrio thiophilus*, a gram-negative, non-sporulating bacterium residing and thriving in water sediments under anaerobic and microaerophilic conditions and that reduces sulfur and nitrate ^34^. We discovered that the flagellar surface of *G. thiophilus* carries significantly higher negative charge, suggesting a potential mechanism for its adhesion to positively charged minerals. Moreover, we observed remarkable diversity in the outer domains of the three newly elucidated structures in both size and fold. The only similarity was that the D2 domain of *S. maltophilia* and the D3 domain of *G. thiophilus* both exhibit an immunoglobulin-like (Ig-like) fold topology^35^. This discovery prompted us to conduct a comprehensive review of annotated bacterial flagellins using AlphaFold predications, classifying the predicted flagellin outer domains into 682 structural clusters. Our results indicated that near half of the flagellin sequences with outer domains contain at least one Ig-like domain, suggesting that the Ig-like domain is ubiquitous in flagellin proteins. Additionally, our analysis provided a profile of the most frequently observed outer domains and determined whether their structures have been experimentally resolved. Ranked by the abundance of outer domain architectures, prior cryo-EM studies have documented outer domain structures in clusters #2, 3, 5, 20, and 28, while X-ray studies have explored outer domains in clusters #1, 2, 5, 6, 10, and 17. The structures we reported belong to clusters #1, 27, and 41, addressing a significant gap in knowledge and highlighting the existence of numerous other fascinating clusters yet to be discovered in structural studies.

## Results

### Cryo-EM structures of three flagellar filaments with outer domains

Cryo-EM was utilized to determine the structures of three flagellar filaments from *C. gilardii*, *S. maltophilia*, and *G. thiophilus*. The established helical symmetry of canonical flagellar D0/D1 domains, with a helical rise of ∼ 5.0 Å and a twist of ∼ 65.45 degrees, was confirmed by examining the power spectra. When this symmetry was applied to 320-pixel box particles, it led to high-resolution 3D reconstructions: *C. gilardii* D0/D1 domains at 3.1 Å, *S. maltophilia* D0-D2 domains at 3.2 Å, and *G. thiophilus* D0-D3 domains at 3.4 Å. This indicates that the D2 domain of C. gilardii and D4/D5 domains of *G. thiophilus* diverge from canonical flagellar symmetry, featuring additional interfaces within the outer domain region (**Fig. 1**). In order to generate 3D reconstructions for supercoiled flagellar filament, particles were re-extracted using a larger 640- pixel box, followed by subsequent inspection of power spectra to identify a layer-line with meridional intensity that does not exist in the canonical flagellar symmetry. A weak meridional layer-line near 1/(110 Å) was observed in *G. thiophilus* flagella, suggesting a repeating feature every 22 flagellin molecules, similar to the flagellar filament in *P. aeruginosa* PAO1^16,18^. Additionally, previously described intermediate layer-lines^18,36–38^ were seen in both power spectra of *C. gilardii* and *G. thiophilus* flagella, indicating non-helical perturbations were present in the structure. As described in Supplemental Figure 1, the reconstruction of the supercoiled filament was executed using helical refinement applying the respective outer domain symmetry and post-processing with homogeneous and local refinements to relax the helical constraints. The 3D reconstructions of the supercoiled filaments for *C. gilardii*, *S. maltophilia*, and *G. thiophilus* achieved final, near-atomic resolutions of 3.4 Å, 3.3 Å, and 4.1 Å, respectively, as determined by map:map FSC (Supplemental Figure 2).

**Fig. 1.**
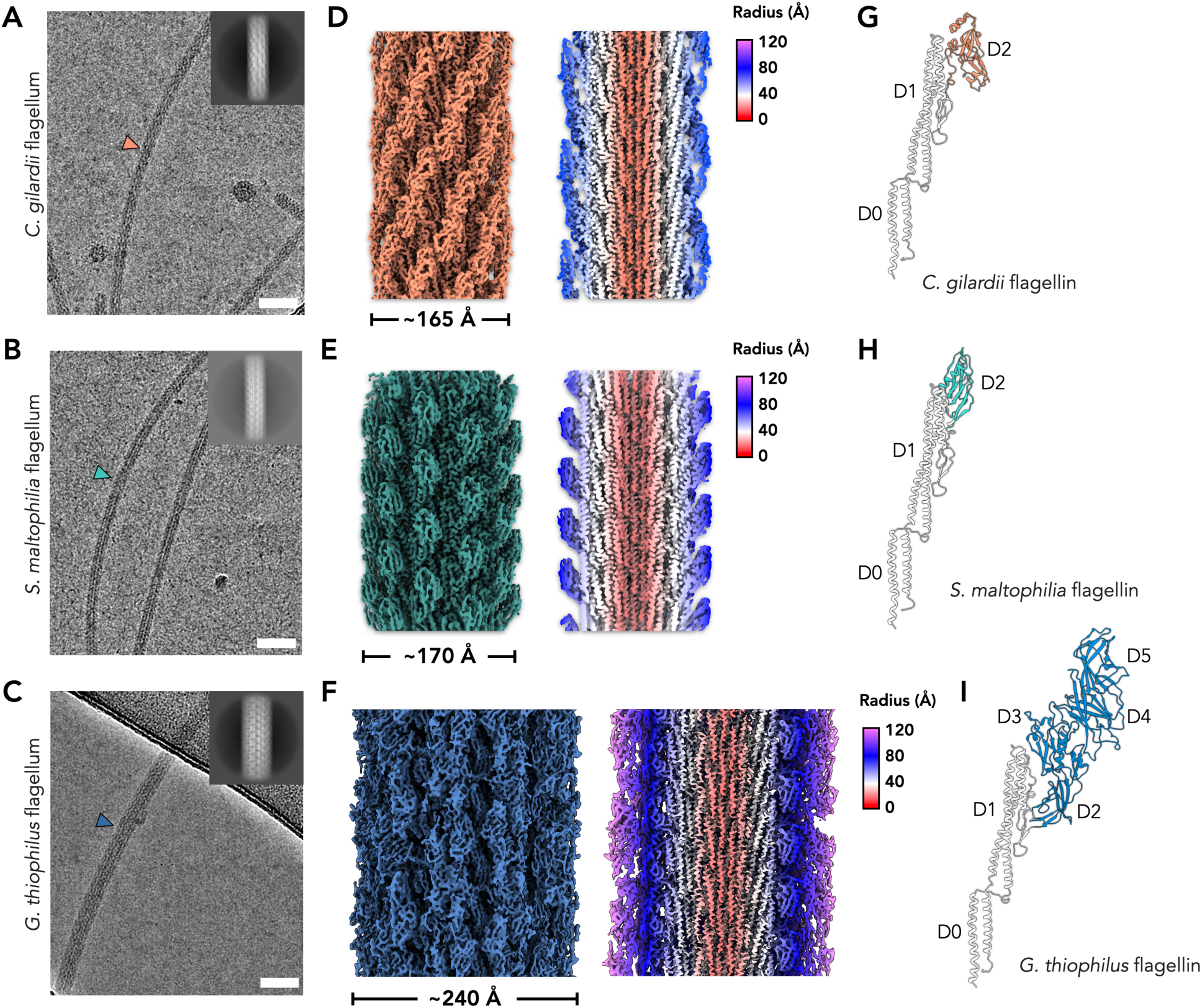
Cryo-EM of flagellar filaments from *C. gilardii*, *S. maltophilia*, and *G. thiophilus*. Representative cryo-electron micrographs of the *C. gilardii* flagella (**A**) *S. maltophilia* flagella (**B**) and *G. thiophilus* flagella (**C**). Scale bar, 50 nm in **A**-**C**. Orange arrowhead points to the *C. gilardii* flagellum; green arrowheads point to the *S. maltophilia* flagellum; blue arrowhead points to the *G. thiophilus* flagellum. 2D averages of all three flagella are shown in the top right corner. Cryo-EM reconstructions of the *C. gilardii* flagellum at 3.4 Å resolution (**D**), the *S. maltophilia* flagellum at 3.3 Å resolution (**E**), and the *G. thiophilus* flagellum at 4.1 Å resolution (**F**). Thin sections parallel to the helical axis of the flagella are shown on the right of the 3D reconstruction, colored by the radius. (**G-I**) The single flagellin structures modeled into the cryo-EM densities are displayed on the right, with their respective domain names indicated. The conserved D0/D1 domains are shown in grey. The outer domains of C. gilardii, S. maltophilia, and G. thiophilus are consistently colored in accordance with the 3D reconstructions.

The diameters of *C. gilardii* and *S. maltophilia* flagellar filaments were similar, approximately 165-170 Å, in contrast to the notably wider *G. thiophilus* filament, which measured around 240 Å (**Fig. 1A-C**). All three flagellar filaments exhibited a similar D0/D1 domain architecture, which is expected since all bacterial flagella share this homology. Notably, both *C. gilardii* and *S. maltophilia* have a single D2 domain. In *C. gilardii*, the D2 domain dimerizes, creating a screw-like feature on its surface (**Fig. 1D**). Conversely, in *S. maltophilia*, the D2 domain does not form an extra interface with adjacent domains, maintaining the same helical symmetry as the D0/D1 domain. The outer domains D2-D5 in *G. thiophilus* are the largest flagellin outer domains determined by cryo-EM (**Fig. 1F, I**), responsible for the diameter increase to 240 Å. Interestingly, three flagellin sequences are present in the *G. thiophilus* genome, quite similar in size and overall folds predicted by AlphaFold. Upon full length modeling of all three flagellin sequences, the correct flagellin protein sequence was identified, as particular regions in the reconstruction were found that could not be explained by the other two sequences.

### Outer domains in three flagellins share β-strand rich architecture

As expected, the D0/D1 domains among the three flagella filaments share an identical fold, with sequence identities ranging from 42-59%. However, the fold of their outer domains significantly diverge (**Fig. 1G-I**). Remarkably, the fold of the *C. gilardii* D2 domain (**Fig. 2A**) was not observed in known, experimentally determined structures, as indicated by Foldseek^39^ and the DALI^40^ server. Further investigation into sequence-level and AlphaFold prediction similarities revealed that similar outer domains are present in certain species of the *Betaproteobacteria* class, including those in the *Burkholderiales* order, and in species of the *Gammaproteobacteria* class, including those in the orders *Enterobacterales*, *Oceanospirillales*, and *Pseudomonadales*, many of which are opportunistic human and plant pathogens. It’s probable that the species belonging to those groups have been sparsely sampled in the past for structural studies. The *S. maltophilia* D2 domain exhibits an Ig-like fold (**Fig. 2B**), a common two-layer β-sandwich domain found in numerous proteins with diverse functions^35^, including the globular domain in archaeal type IV pili^41–43^. Unsurprisingly and likely due to the ubiquitous nature of the Ig-like domain, similar outer domains were found in many AlphaFold predicted flagellins across a diverse array of bacterial species, from anaerobes to aerobes and marine algae to human microbiota. A similar D2 domain was also experimentally captured in a partial flagellin crystal structure from *Sphingomonas sp.* A1^19^, sharing approximately 28% sequence identity between the D2 domains. The *G. thiophilus* flagellum, notably the first flagellar structure from bacteria growing anaerobically, presents a large outer domain architecture with four domains D2-D5 (**Fig. 2C**). All four outer domains are rich in β-strands: the fold of D2 domain is mainly found in flagellar hook-associated proteins; the D3 domain is a variant of the Ig-like fold; D4 and D5 are β-sandwich domains similar to the Pfam^44^ DUF992 (domain of unknown function). Similar flagellar outer domains are mostly identified in bacteria from environments like lake or ocean sediment, soil, and wastewater, across the classes *Chrysiogenetes*, *Clostridia*, *Deferribacteres*, and *Synergistia*.

**Fig. 2.**
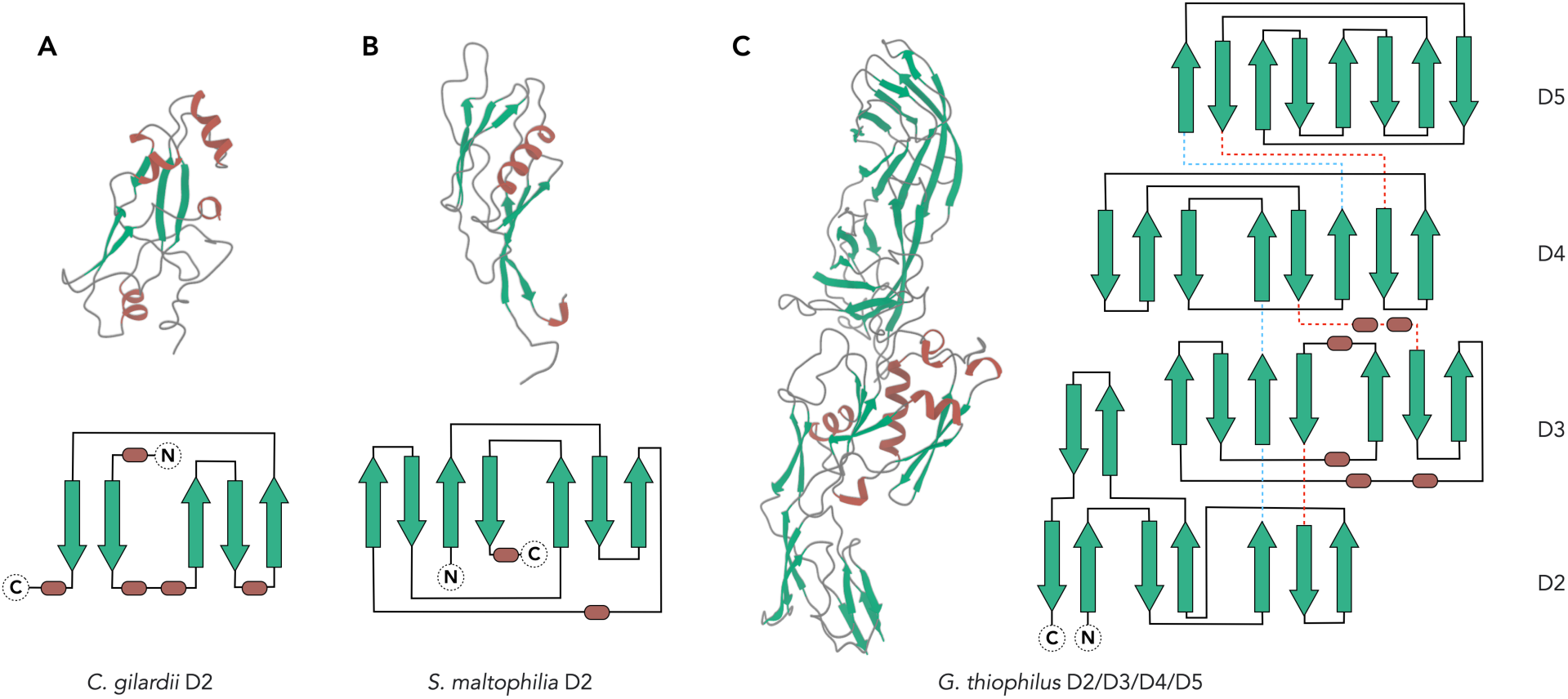
The fold of flagellar outer domains from *C. gilardii*, *S. maltophilia*, and *G. thiophilus*. Cryo-EM structures of the flagellar outer domains from *C. gilardii* (**A**), *S. maltophilia* (**B**), *and G. thiophilus* (**C**) were presented, with all α-helices colored brown and β-sheets colored green. These domains are also drawn in schematic representation, with β-sheets shown as green arrows, α-helices as brown cylinders, and loops as dark grey lines. When multiple domains are present, they are connected by blue dashed lines near the N-terminus and red dashed lines near the C-terminus.

### Dimeric outer domain interactions in *C. gilardii* and *G. thiophilus* flagella

Next, we explored how the outer domains interact along the flagellar filaments. Interestingly, the outer domain of *S. maltophilia* maintains the same D0/D1 symmetry and does not form additional polymeric contacts with adjacent subunits. In contrast, the *C. gilardii* and *G. thiophilus*flagella exhibit partial or entire outer domain dimerization, thereby disrupting the D0/D1 symmetry. The dimeric interface happens at the radius of ∼80 Å in *C. gilardii* and ∼120 Å in *G. thiophilus* (**Fig. 3A**). Given the canonical D0/D1 symmetry, adjacent flagellins along the 11-start have minimal twist (∼65.45 degrees multiplied by 11), rendering the protofilament nearly parallel to the helical axis. In both *C. gilardii* and *G. thiophilus*, the outer domains — D2 in *C. gilardii* and D4/D5 in *G. thiophilus* — protrude from the 11-start and adopt either an “up” or “down” conformation.

**Fig. 3.**
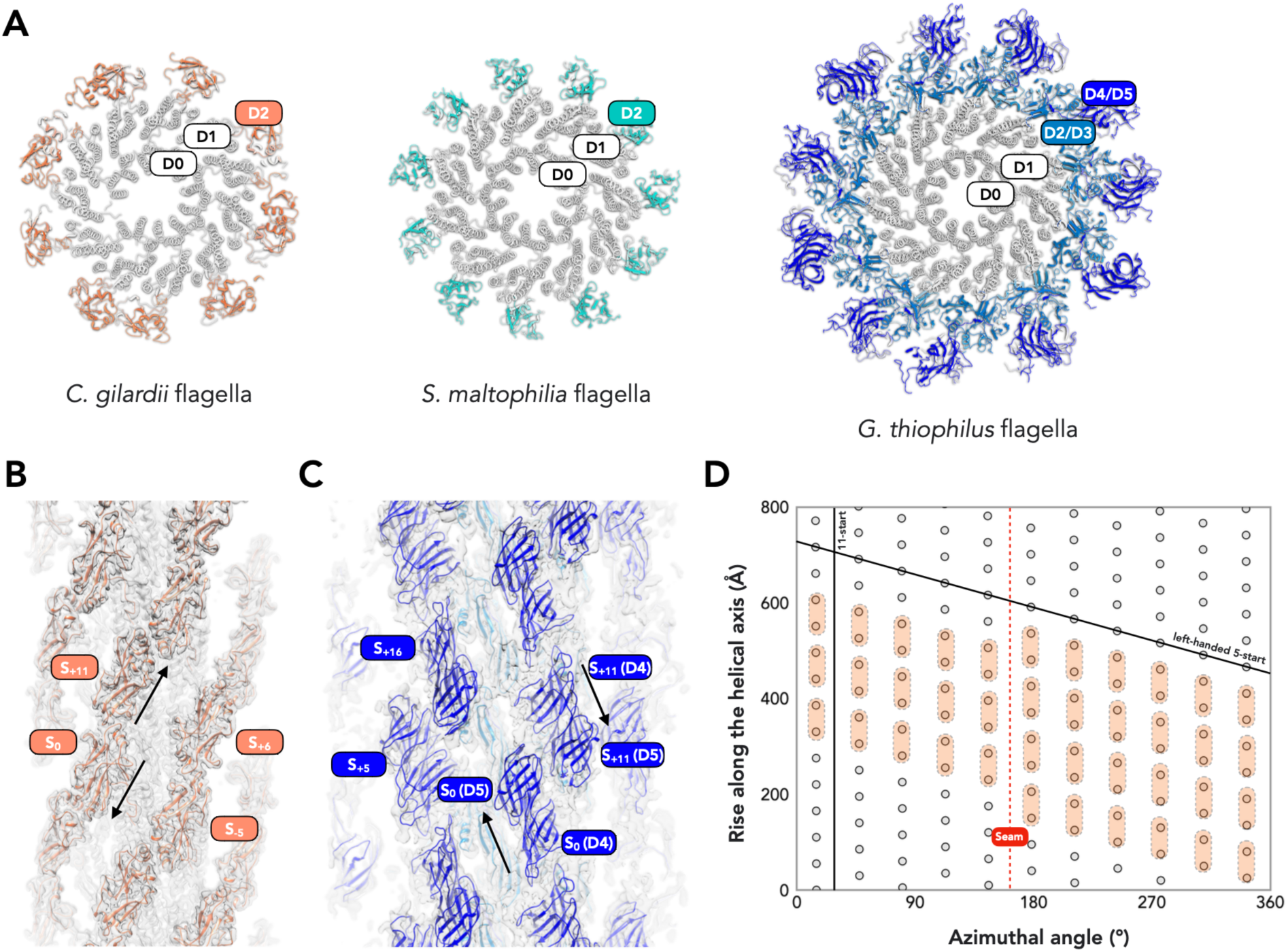
Outer domain arrangements on flagellar surface. (**A**) The top view of three flagellar filaments with the conserved D0/D1 domains in grey, the D2 domain of *C. gilardii* flagellum in orange, the D2 domain of *S. maltophilia* flagellum in green, the D2/D3 domains of *G. thiophilus* flagellum in light blue, and the D4/D5 domains of *G. thiophilus* flagellum in dark blue. (**B**) The D2 domain dimerization in *C. gilardii* is presented within the cryo-EM map density. (**C**) The D4/D5 domain dimerization in *G. thiophilus*, with the model shown within cryo-EM map density. (**D**) The helical net of the flagellar filament is illustrated using the convention that the surface is unrolled, providing a view from the outside. Grey dots indicate the approximate symmetry present in the D0/D1 domains of all flagellar filaments. The conventional 11-start and left-handed 5-start, originating from the D0/D1 domain, are indicated with black lines. The dimer interface that occurs in the flagellar outer domains of both *C. gilardii* and *G. thiophilus* is illustrated with transparent orange stadium shapes. The seam arising from this dimerization packed along the D0/D1 5-start is indicated by a red dashed line.

This dimeric interface, formed between the “up” and “down” conformations of the outer domains, generates a screw-like feature in the *C. gilardii* flagellum, where the surface area buried between subunits is approximately 270 Å^2^ (**Fig. 3B**). Conversely, the *G. thiophilus* outer domain lacks a distinct screw-like feature, with the dimeric interface between D5 domains of subunit S_0_ and subunit S_+11_ enclosing a buried interface of about 220 Å^2^. The interaction area between D4 domains along the 11-start protofilament is relatively small, at around 80 Å^2^ as estimated by PDB PISA^45^ (**Fig. 3C**). Notably, the D2/D3 domains in *G. thiophilus*, located at a smaller cylinder radius of about 80 Å, do not dimerize, preserving the approximate symmetry seen in D0/D1.

The dimerization interface occurs between subunits on the same 11-start protofilament, with dimer packing extending along the left-handed 5-start protofilament. Thus, a seam occurs between two 11-start protofilaments (**Fig. 3D**), similar to flagella previously studied in *P. aeruginosa* PAO1^16,18^. This seam, reminiscent of those observed in microtubule filaments, interrupts the continuity along the dimer extension, which is the left-handed 5-start of D0/D1 domains (**Fig. 3D**). Although such structures with a seam were termed “non-helical” in the past, they can be viewed as helical polymers with a large asymmetric unit comprising 22 flagellin subunits while ignoring flagellar supercoiling. This larger ASU reconstruction strategy failed to reach high resolution seven years ago in the reconstruction of *P. aeruginosa* PAO1 flagella data^18^ recorded using a Falcon II camera in integrating mode and processed with legacy software SPIDER^46^. However, advancements in cryo-EM now make this strategy possible (Supplemental Figure 1) using a Gatan K3 camera in counting mode, processed in CryoSPARC^47^.

### The flagellar surface of G. thiophilus is negatively charged

The flagellar outer domains of *C. gilardii* and *S. maltophilia* are significantly smaller compared to the outer domains in *G. thiophilus*, thereby failing to completely shield the inner D0/D1 domain from solvent exposure. In contrast, *G. thiophilus* possesses a two-layered outer domain, comprised of the D2/D3 middle layer and the D4/D5 outer layer, which effectively encapsulates the D0/D1 domain from solvent access. This observation prompted us to investigate whether such coverage impacts the surface charge by estimating the coulombic electrostatic potential for the flagellar surfaces of these three species. Remarkably, the surface of the *G. thiophilus* flagellum is significantly more negatively charged than those of the other two species, attributed to a higher presence of aspartic acid and glutamic acid as surface residues (Supplemental Figure 3A); analysis of amino acid composition suggested the same conclusion. When examining the D0/D1 domains, the amino acid distribution among the three flagella was nearly identical. However, a comparison of the outer domains revealed that the *C. gilardii* flagellum, while having an overall amino acid composition similar to *S. maltophilia*’s, possesses slightly more negatively charged residues, resulting in a mildly negative surface charge. On the other hand, the outer domain of the *G. thiophilus* flagellum displays a notable difference, containing three times as many negatively charged residues compared to positively charged ones. Even though the *G. thiophilus* outer domain is approximately 3.5 to 4 times larger than those of the other two flagella, it has 6 to 7 times more negatively charged residues, resulting in a predominantly negatively charged surface that fully covers the D0/D1 domain (Supplemental Figure 3B-C).

### The Ig-like fold is ubiquitous in the bacterial flagellin outer domain universe

Flagellar outer domains are known to vary greatly in protein sequence and length^48^, which is consistent with our findings. We observed that two of the three flagellins we report contain an Ig-like domain, which have been noted in several other previously reported flagellin structures^16,18–20,22^. This prompted us to question whether flagellin outer domains possess certain folds/domains more commonly than others. The advent of AlphaFold^28^ has enabled this type of computational analysis. To date, all AlphaFold predictions have been clustered based on fold similarity^49^. These results, unfortunately, were biased by D0/D1 alignment and could not be directly applied to our study, which seeks domain clustering results independent of D0/D1 domains. Therefore, we downloaded all bacterial flagellin AlphaFold predictions for analysis, trimming the D0/D1 domains (263 residues) based on prior knowledge of experimental flagellin structures. Interestingly, about 60% of flagellins are shorter than 350 residues (**Fig. 4A-B**), containing either only the D0/D1 domain, such as the *Bacillus subtilis* flagellum^18^, or the D0/D1 domain of *Borreliella burgdorferi* flaA^50^, which contains an extra small domain/disordered loop (**Fig. 4B**). Thus, we set a cutoff at 350 residues for substantial outer domains analysis, creating a library of 16,948 bacterial flagellin outer domain predictions.

**Fig. 4.**
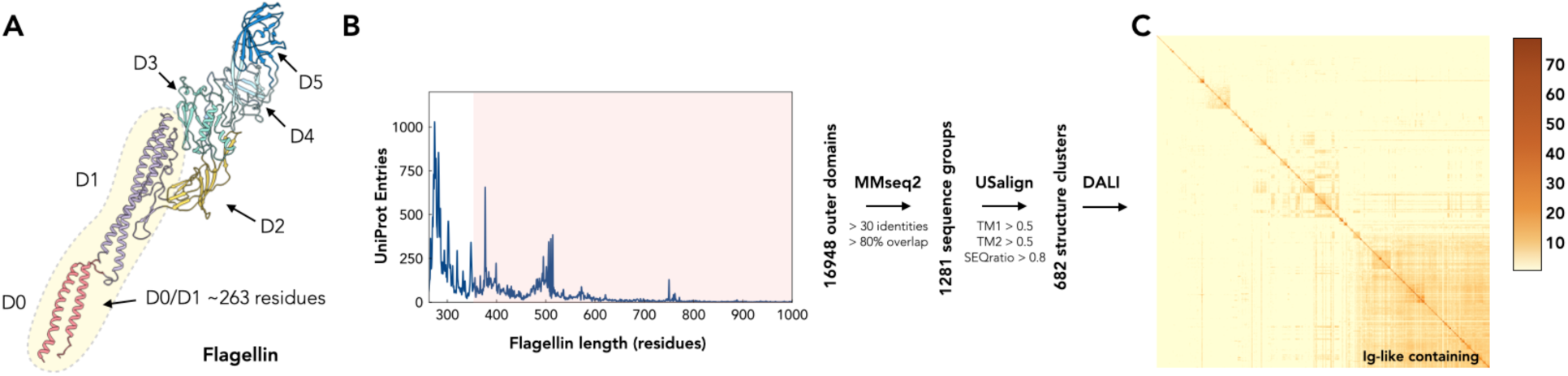
Analysis of AlphaFold predictions of bacterial flagellin outer domains. (**A**) The typical architecture of flagellin including outer domains. The D0 and D1 domains, which have 263 or a bit more residues, are colored in red and purple, respectively. The other outer domains, extending from inner to outer diameter, are sequentially named D2, D3, D4, and so on. (**B**) This plot represents the number of UniProt entries corresponding to various lengths of flagellin proteins. Flagellin AlphaFold predictions equal and longer than 350 residues are selected for subsequent multistep analysis, as described in the method section. 682 structural clusters were generated (**C**) The matrix visualizes a DALI all-to-all analysis of 682 structural cluster representatives. It is constructed using pairwise DALI Z-scores, with the corresponding Z-score color scale presented on the right. The most prominent cluster, located at the bottom right of the matrix, contains representatives with Ig-like domains.

For initial clustering, we employed a protein sequence-based approach, similar to, yet more aggressive than AFDB clustering, by grouping similar protein sequences with MMseqs2^51^ at a 30% sequence identity and 80% overlap cutoff. This method categorized the 16,948 predictions into 1,281 sequence-based groups. We noticed many flagellin outer domains exhibited a variable number of domains present. Taking this into account, we next applied a more conserved rigid-body alignment strategy by utilizing USalign^52^ combined with graph-based community detection clustering via the walktrap^53^ algorithm to further group entries with similar lengths and structures. This step organized 1,281 sequence-based groups into 682 structural clusters. Each cluster contained a unique “representative” flagellin, distinct from others at the sequence or rigid-structure level (**Fig. 4B**). Lastly, we examined whether prevalent domain folds were maintained within this library of 682 structural clusters. We used DALI, a tool for analyzing deep phylogenetic relationships and protein homology, known for its sensitivity to fold similarities, to perform an all-to-all analysis^40^. Strikingly, this analysis revealed significant clustering, as shown in the Z-score heatmap (**Fig. 4C**), with about 40% of the representatives carrying at least one Ig-like domain, comprising nearly half of the 16,948 AlphaFold predictions.

### Diverse outer domains remain to be studied in bacterial flagella

Lastly, we asked what the most abundant folds/domains are in known bacterial flagellar outer domains and how they are related. To accomplish this, using the same community-based detection strategy for the USalign clustering discussed above, we grouped the remaining representatives using a DALI Z-score greater than 10 as the cutoff for significance. We listed the most populated structural clusters, the number of predictions within those clusters, AlphaFold predictions of the representatives, and the known experimental structures belonging to a given cluster (**Fig. 5A**). Not surprisingly, the largest cluster in the center is comprised of representatives that have at least one Ig-like domain within the outer domains. Among 682 structural clusters, five out of the six most populated clusters contain one or more Ig-like domains: cluster #1 with 2,304 entries, including *S. maltophilia* flagellin; cluster #2 with 1,838 entries, featuring two Ig-like domains as observed in *P. aeruginosa* PAO1^16,18^; cluster #3 with 1,094 entries, including another variant of two Ig-like domains, as in *Campylobacter jejuni* flagellin^20^; cluster #4 with 1,094 entries, possessing one Ig-like and one β-barrel domain; and cluster #6, showcasing a single Ig-like domain variant previously identified in a different strain of *P. aeruginosa*^22^ (**Fig. 5B**).

**Fig. 5.**
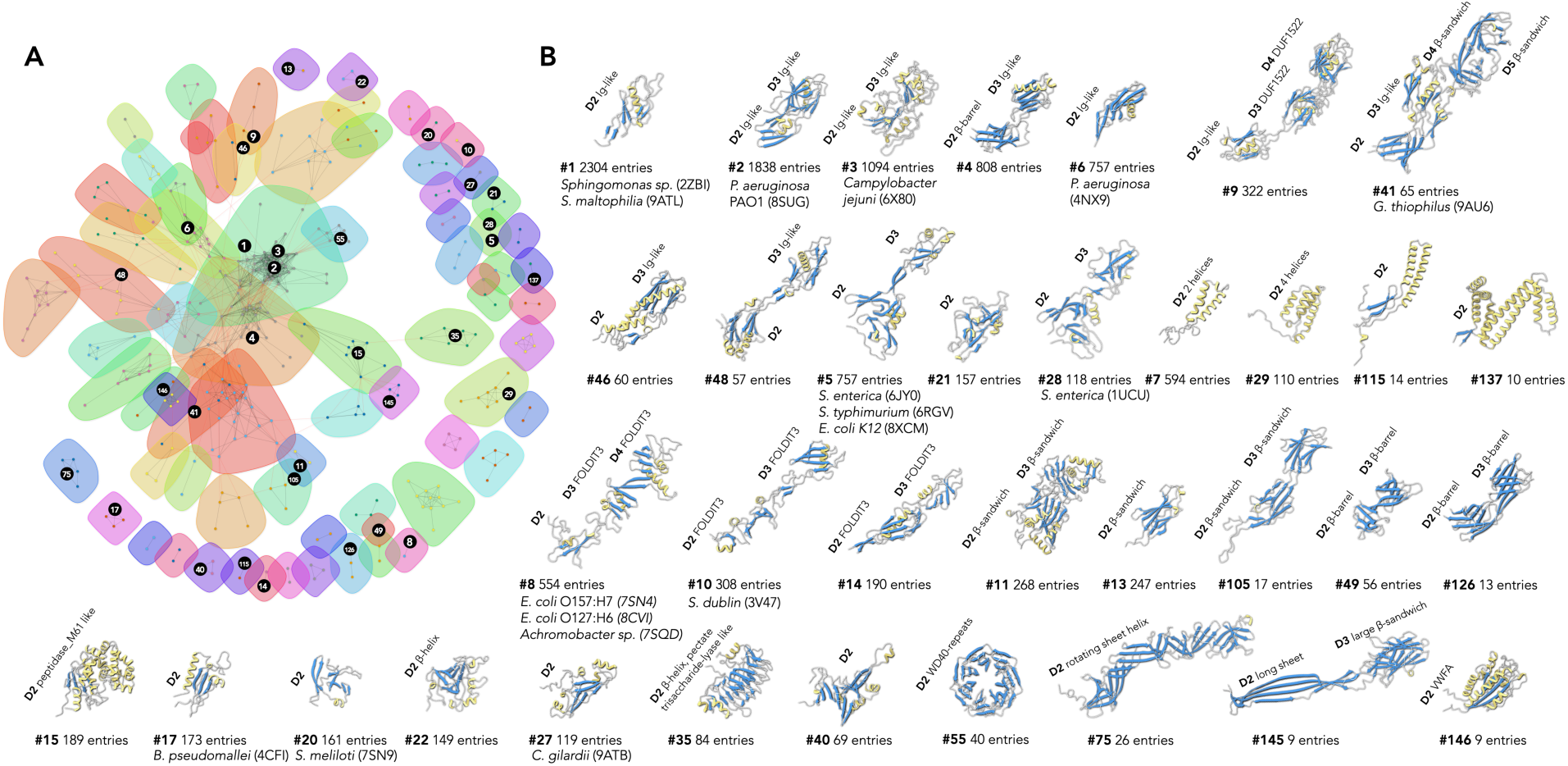
Populated flagellin outer domains in the bacterial domain. (**A**) The relationships among 682 structural clusters are depicted using the Fruchterman-Reingold algorithm, a force-directed layout algorithm from the R/igraph package. The communities, indicated with transparent color circles, were detected using the walktrap algorithm with a step size 6. Additionally, some most populated clusters, as illustrated in (**B**), are highlighted with black circles. (**B**) AlphaFold predictions or experimentally determined structures from populous structural clusters are organized according to the population size of the first representative in a given community. Within these models, all α-helices are colored yellow, β-sheets blue, and loops grey. Known domain folds are labeled. For clusters with available cryo-EM or X-ray structures, the corresponding species and PDB IDs are provided.

Apart from the Ig-like domain, we observed various other widespread folds in flagellin outer domains. For instance, several flagella structures in similar clusters like #5, #21, and #28 have been reported, including *Salmonella enterica*^21^, *Salmonella typhimurium*^10^, and *Escherichia coli* K12^17^. Structural clusters mainly comprising α-helices, such as #7, #29, #115, etc., were also identified. Interestingly, no experimental structure has been documented in these α-helix rich clusters yet. Other clusters having experimental structures include cluster #8, featuring *E. coli* O157:H7, *E. coli* O127:H6, and *Achromobacter spp*. flagella^17^, and cluster #10, containing *Salmonella dublin*^23^ flagellum. These outer domains typically exhibit “skinny” architectures containing domain folds similar to the *de novo* designed domain named foldit3^54^. Other less abundant, but intriguing, clusters were also observed; for instance, the outer domain in cluster #15 resembles the peptidase-M61 catalytic domain^55^; the fold in cluster #35 is similar to the pectate trisaccharide-lyase domain^56^; and the WD40-repeats in cluster #55, more commonly found in eukaryotes, is often thought to serve as a rigid scaffold for protein interactions^57^. Two of the three flagellar structures reported in this study belong to previously unexplored outer domain clusters: *C. gilardii* (cluster #27) and *G. thiophilus* (cluster #41). Although these newly discovered families are not the most abundant in our analysis, there may be a bias introduced by commonly studied bacterial species that are more redundantly sampled in the database. Overall, this analysis suggests that many more folds exist in flagellar filaments that have yet to be sampled, and their functions remain to be discovered.

## Discussion

The field of structural biology has rapidly advanced due to recent developments in cryo-EM^58,59^ and structure prediction^28^ techniques. These developments, when combined with experimental data, now enable the performance of large-scale^29^ and robust structural analyses at the protein fold level. In this study, we determined the cryo-EM structures of three bacterial flagellar filaments at near-atomic resolution. These structures encompass both the conserved D0/D1 regions and the outer domains, providing a general 3D reconstruction strategy for wild-type supercoiled bacterial flagella. Among our findings, we identified novel folds not previously observed in other bacterial flagella. Of particular interest, was the prevalent Ig-like fold within the known flagellar outer domains. Our in-depth analysis of bacterial flagellar outer domains confirmed the widespread presence of the Ig-like domain; it is found in nearly half of all known flagellin proteins that are longer than 350 residues. Additionally, we discovered other interesting flagellar outer domains, which could spark future research to study flagellar filaments in unexplored species.

From the perspective of bacterial “economics”, flagellin emerges as one of the most energetically demanding proteins for bacteria to produce, requiring an estimated 20,000 copies per flagellum. This high energy investment in flagella production was clearly demonstrated over 30 years ago in studies when bacteria were grown in stirred liquid cultures where motility provides no advantage but incurs additional energy expenditure. Bacterial cells lost their flagellar filaments resulting from spontaneous mutations in flagellar genes within just 10 days, highlighting an evolutionary advantage for bacteria that conserve energy by not producing flagella^60^. Recent estimates suggest that the energy cost of synthesizing flagellar filaments accounts for 0.5% to 40% of the total energy budget across different species^61^. The presence of outer domains in flagella, especially in species like *G. thiophilus*, where the flagellin size can reach up to 793 residues—approximately triple the size of a flagellin composed solely of the D0/D1 domain—underscores the hypothesis that these elaborate structures must confer substantial benefits to justify their high energy costs.

While the conserved D0/D1 domain is crucial for swimming motility, the necessity of the outer domains for this function has been questioned. In *S. typhimurium*, it has been shown that flagella lacking an outer domain remain motile, suggesting that this domain is not required for swimming motility^15^. Yet, studies suggest that the outer domain may enhance motility in several ways. For instance, in *E. coli*, the outer domains have been shown to extend tumbling time, thereby improving navigation efficiency^17^. Similarly in *P. aeruginosa* PAO1, mutations in the outer domain disrupt the formation of supercoiled filaments, adversely affecting motility^16^. Conceptualizing the flagellar filament as a propeller, the outer domain could enhance the motility by increasing the flagellar diameter and adjusting the angle of attack, potentially aiding movement in high-viscosity environments. Beyond motility, outer domains may serve diverse functions across different bacterial species. For example, the outer domains in cluster #15 resemble glycyl aminopeptidase, suggesting a role in digesting extracellular proteins to facilitate the uptake of essential amino acids^62^. In the case of *G. thiophilus* (cluster #41), which thrives in anoxic or microaerophilic environments, and has the largest flagellin structure identified, its flagellum features an unusually negatively charged surface. This characteristic might regulate motility modes in various environments. For instance, in the presence of positively charged surfaces like minerals, iron oxides, and most metal oxides, the flagellum may adhere more readily, leading bacteria to detach their flagella and colonize the surface. Conversely, in environments that are more negatively charged and devoid of minerals, the flagellum could experience reduced resistance, allowing for smoother rotation and enhanced motility.

It has been proposed that the outer domain may potentially offer antigenic variability, aiding in the evasion of immune surveillance^36^. Interestingly, in the well-studied Toll-like receptor 5 (TLR5) pathway, previous structural studies suggested flagellin binding induced TLR5 dimerization^23,63^. In its monomeric form, the flagellin outer domain, in species such as *C. gilardii* and *S. maltophilia*, is not large enough protect the D1 conserved site. However, in its filament form, outer domains could potentially help fight against adaptive immunity. Intriguingly, *Helicobacter pylori* and *Campylobacter jejuni* possess mutations in this TLR5 binding D1 region, thereby possibly evading the TLR5 immune response^64,65^. Furthermore, *H. pylori* flagella exhibits an additional layer of complexity with its flagella encased in a membrane sheath that completely covers the supercoiled structure^66^. This sheath is present even though the flagella retain their outer domains. Those observations suggest functions of the outer domain in *H. pylori* and *C. jejuni* are beyond merely shielding the D0/D1 domain, and probably essential for the filament integrity.

Several studies have shown that flagellar outer domains can undergo additional surface structure organization, disrupting the D0/D1 symmetry, resulting in the formation of structures such as dimers, dimers with seams, tetramers, and tetramers with seams^16–18^. Notably, despite the similarity in flagellar outer domain architectures between *E. coli* O157:H7 and *E. coli* O127:H6, which both belong to cluster #8, they exhibit distinct surface organization patterns: O157:H7 forms dimers without a seam, whereas O127:H6 assembles into tetramers with a seam. On the other hand, dimers with seams are commonly observed in flagellar outer domains, with known cryo-EM structures identified in bacteria like *P. aeruginosa* PAO1 (cluster #2), *C. gilardii* (cluster #27), and *G. thiophilus* (cluster #41). This suggests that such surface organization reflects a strategy for optimized spatial packing, influenced more by the size and surface properties of the outer domains than by the protein fold or specific function. From the helical net (Fig. 3C), it becomes apparent that the linker connecting D1 and D2 domains in subunit S_0_ is positioned close to S_5_, S_6_, and S_11_. Should dimerization occur between S_0_ and S_11_, with the resulting protofilament extending along the left-handed 5-start (all the numbered start refers to the classic D0/D1 symmetry), a seam will form (Fig. 3C). Conversely, when dimerization occurs between S_0_ and S_5_, extending along the right-handed 6-start, as seen in *S. meliloti*^17^, no seam is formed. Thus, it is reasonable to anticipate the discovery of other packing arrangements, such as trimerization or pentamerization, within the flagellar outer domain in the future. Such diversity in packing could further reveal the adaptability and complexity of flagellar structure and function.

Bacteria often harbor multiple flagellin genes, such as those seen in *G. thiophilus*, suggesting the potential for diverse functionality fulfilled by different flagellin variants, which are produced in response to environmental needs. For instance, certain bacteria such as *Azospirillum lipoferum*^67^ and *Shewanella piezotolerans*^68^, produce polar flagella for swimming in liquid environments and multiple lateral flagella for swarming on surfaces, demonstrating a sophisticated level of adaptive behavior. A similar phenomenon has recently been observed in the archaeal domain, where *Sulfolobales islandicus* REY15A can utilize the same pilin protein to generate two distinct type IV pili structures by modulating its Ig-like outer domain in response to different environmental needs^41^. This underscores a broader principle of microbial adaptability and the strategic regulation of extracellular filament secretion. Clearly, despite these advances, our understanding of the regulatory mechanisms governing flagellin gene expression, extracellular filament secretion and the specific roles of novel outer domains remains incomplete. Future research dedicated to unraveling these aspects promises to deepen our insights into microbial motility and interaction with their environments.

## Material and Methods

### *C. gilardii* flagellar filament preparation

*C. gilardii* cells were grown aerobically at 37 °C for 48 hours in chemical defined medium^69^ in 5% CO_2_ incubator. For isolation of filaments, cells were spin down at 7,000 rpm (9,120 x g, Beckman JLA, 12.500 rotor). Pellet was suspended with 4 ml phosphate-buffered saline (PBS) buffer pH 7.2, and filaments were sheared off from the cell using a homogenizer at 10,000 rpm for 10 min. Cells and debris were removed by centrifugation at 10,000 x g for 10 min. After this, filaments were pelleted from the supernatant by ultra-centrifugation (Beckman 50.3 Ti ultra-rotor, 35,000 rpm) at 4°C for 1.5 h, and subsequently resuspended with 150 µl PBS buffer. DNase I (NEB) were added into the sample to remove possible extracellular DNA fibers.

### *S. maltophilia* flagellar filament preparation

*S. maltophilia* cells were grown aerobically at 37 °C for 48 hours in Tryptic Soy Broth (TSB, BD^TM^) medium in 5% CO_2_ incubator. The resultant pellet was resuspended in 4 mL of PBS buffer, and the cell suspension was put under a homogenizer for 10 min to shear off the extracellular filaments as described above. The cells were then removed by centrifugation at 10,000 x g for 10 min. The supernatant was collected and the filaments were pelleted by ultracentrifugation (Beckman 50.3 Ti ultra-rotor, 35,000 rpm, 1.5 h, 4°C). After the run, the supernatant was removed, and the pellet was resuspended in 150 µL of PBS. DNase I (NEB) were added into the sample to remove possible extracellular DNA fibers.

### *G. thiophilus* flagellar filament preparation

*G. thiophilus* cells were grown anaerobically at 30 °C in anaerobic freshwater medium 503 (DSMZ) in a volume of 10 ml. After this, cells were vortexed for 30 min to mechanically shear off the extracellular filaments, and cells were subsequently removed by centrifugation at 10,000 g for 10 min. The supernatant was collected, and filaments were further enriched by an overnight 20% ammonium sulfate precipitation at 4 °C. The resultant flagellar filament pellet was collected and resuspended with 150 µL 100 mM ethanolamine buffer pH 10.5, incubated with Dnase I (NEB) prior to plunge freezing.

### Cryo-EM conditions and image processing

The flagellar filament sample (4.5 μl) was applied to glow-discharged lacey carbon grids and then plunge-frozen using an EM GP2 Plunge Freezer (Leica). The cryo-EM micrographs were collected on a 300 keV Titan Krios with a K3 camera at 1.11 Å per pixel and a total dose of 50 e^-^/Å^2^. The cryo-EM workflow was initiated with patch motion corrections and CTF estimations in cryoSPARC^47,70,71^. Following this, an automated picking of particle segments was conducted using the ‘Filament Tracer’ function with a shift of 10 pixel between adjacent boxes. All auto-picked particles were subsequently 2D classified with multiple rounds, and all particles in bad 2D averages were removed. Next, the possible helical symmetries were calculated from averaged power spectra generated from the raw particles (640-pixel box)^72,73^. The identified parameters consistent with canonical flagellar filaments were then applied in 3D helical refinement to generate a high-resolution map of the D0/D1 domains. Subsequently, the power spectra’s meridian area was carefully examined to detect layer-lines that might indicate interfaces of additional outer domains. Upon the identification of such layer-lines, a re-extraction of particles was performed, applying a shift slightly greater than the periodicity observed in the power spectra to avoid particle duplications. After that, 3D reconstruction was performed using “Helical Refinement” first, “Homogenous Refinement” next, then “Local Refine”, Local CTF refinement, and another round of “Local Refine” using CTF-refined particles (Supplemental Figure 1). The resolution of each reconstruction was estimated by Map:Map FSC, Model:Map FSC, and d_99_^74^. Maps used in the resolution estimation were sharpened using Local Filter available in cryoSPARC. The statistics are listed in Table 1.

**Table 1.** Cryo-EM and Refinement Statistics of flagellar filaments.

| Parameter | *C. gilardii* flagellum | *S. maltophilia* flagellum | *G. thiophilus* flagellum |
| --- | --- | --- | --- |
| **Data collection and processing** |  |  |  |
| Voltage (kV) | 300 | 300 | 300 |
| Electron exposure (e<sup>-</sup> Å<sup>-2</sup>) | 50 | 50 | 50 |
| Pixel size (Å) | 1.11 | 1.11 | 1.11 |
| Particle images (n) | 223,582 | 275,407 | 21,618 |
| Shift (pixel) | 100 | 10 | 100 |
| **Approximate Helical symmetry before symmetry relaxation** |  |  |  |
| Point group | C1 | C1 | C1 |
| Helical rise (Å) | 110.25 | 5.03 | 111.12 |
| Helical twist (°) | -0.61 | 65.40 | -0.30 |
| **Map resolution (Å)** |  |  |  |
| Map:map FSC (0.143) | 3.4 | 3.3 | 4.1 |
| Model:map FSC (0.5) | 3.6 | 3.5 | 4.5 |
| d<sub>99</sub> | 3.8 | 3.6 | 4.5 |
| **Refinement and Model validation** |  |  |  |
| Ramachandran Favored | 95.8 | 94.1 | 93.0 |
| Ramachandran Outliers | 0.0 | 0.0 | 0.4 |
| Real Space CC | 0.88 | 0.89 | 0.76 |
| Clashscore | 5.7 | 6.0 | 9.5 |
| Bonds RMSD, length (Å) | 0.008 | 0.005 | 0.010 |
| Bonds RMSD, angles (°) | 0.787 | 0.633 | 0.706 |
| **Deposition ID** |  |  |  |
| PDB (model) | 9ATB | 9ATL | 9AU6 |
| EMDB (map) | EMD-43829 | EMD-43830 | EMD-43868 |

### Model building of flagellar filaments

The first step in model building is to identify the correct flagellin protein from the experimental cryo-EM map, especially when there are multiple candidates. This was the case with the G. thiophilus flagellar sample, which had three similar flagellin sequences in its genome: A0A3R5UZF4, A0A3R5UW90, and A0A3R5V2W1. To address this, full-length modeling was carried out for each sequence until a region in the map was encountered that the sequences could not account for (see Supplemental Figure 4). A0A3R5UZF4 was identified as the matching sequence through meticulous examination, fully agreeing with the cryo-EM density.

Regarding the modeling, the AlphaFold predicted structure of a single flagellin subunit was initially docked into the cryo-EM map. Domains (for instance, D0 to D5 in *G. thiophilus* flagella) were docked individually since AlphaFold predictions for domain-domain orientation are frequently imprecise. The subunit was then manually adjusted and refined in Coot^75^. Given that the outer domain forms a dimer in *C. gilardii* and *G. thiophilus* flagella, this procedure was repeated for the other flagellin within the dimer. Following this, the refined single flagellin or flagellin dimers were docked into the other ten protofilaments within the supercoiled flagellar filament map and subjected to real-space refinement via PHENIX^76^. The filament model’s quality was assessed using MolProbity^77^, and the refinement statistics for all three flagellar filaments are detailed in Table 1. Cryo-EM map and model visualization were primarily done in ChimeraX^78^.

### Bioinformatic clustering of flagellin outer domains

First, AlphaFold^28^ predictions of all bacterial proteins annotated in UniProt as “flagellin” were downloaded from the AlphaFold Protein Structure Database^29^. To conduct the analysis in the absence of D0/D1 domains, protein regions corresponding to D0/D1 domains were carefully trimmed from the prediction by excising the first 163 residues and the last 100 residues. This cutoff is selected based on existing experimental flagellin structures. Furthermore, flagellins shorter than 350 amino acids (outer structure shorter than 87 residues) were excluded. This is because the average size of a protein domain is approximately 100 residues, and analysis is based on AlphaFold predictions, which are very meaningful for such a short region. Next, DSSP^79^ was used to estimate the secondary structure components of the remaining models, and models with less than 35% total secondary structures were removed, leaving 16,948 flagellin predictions for clustering analysis.

The initial clustering was performed at the amino acid sequence level using MMseqs2^51^, with the conversion from PDB to FASTA format achieved via Pdb2Fasta. The Mmseqs2 easy-cluster workflow was used based on a minimum of 30% sequence identity, requiring the short sequence to be at least 80% the length of the other sequence. This process reduced the number of 16,948 predictions to 1,281 “sequence groups”, with representatives from each group chosen by MMSeqs2. Further investigation into rigid-body structural similarities among the 1,281 groups was conducted using USalign^52^ pairwise alignments using the semi-non-sequential alignment (sNS or -mm 6) resulting in 819,480 alignments from the 1281 representative structures. An undirected graph was built using the pairwise TM scores, TM1 and TM2 as described in the USalign documentation, and from an alignment length score calculated by taking the length of alignment (Lali in USalign) and dividing it by the total sequence length of the larger outer domain of the two. The cutoffs for edges to be included were TM1 and TM2 greater than 0.5 and the alignment length score greater than 0.8. This was done to include strong alignments yet eliminate edges that corresponded to high TM scores for single domain structures aligning with a similar domain in a multi-domain structure. Using this graph, community detection based clustering was performed using the walktrap algorithm with a step size of 6. From the 1281 sequence groups, 705 were included in the graph with 7115 edges and clustered into 107 communities, the remaining 575 had no significant connections. Merging clustering results from both MMSeqs2 and USalign left us with 682 total clusters encompassing all 16,948 input structures. Clusters were sorted and numbered by member number, largest to smallest. 142 clusters had 10 or more members, 251 clusters had 2 to 9 members, and 274 cluster had only one member, finding no significant structural or sequence similarity through either MMSeq2 or USalign. Within each group, a representative model was chosen for downstream steps by sorting by highest AlphaFold predicted score, pLDDT, and choosing the first structure within five residues of the average sequence length of the cluster. Finally, an all-to-all DALI^40^ alignment was performed among those 682 representatives to identify similar domains within multi-domain flagellar outer structures, as well as similar structures that rigid-body methods failed to align due to different predictions in domain arrangements. A similar strategy to the USalign clustering was used with the cutoff for edge inclusion set at Z-scores above 10. The walktrap algorithm with a step size 6 was again used for community detection.

The graph was drawn using the Fructerman-Reingold algorithm^80^, a force-directed layout algorithm within the R/igraph package.

## Acknowledgments

This research was, in part, supported by the National Cancer Institute’s National Cryo-EM Facility at the Frederick National Laboratory for Cancer Research under contract 75N91019D00024. Electron microscopy screening was carried out in the UAB Cryo-EM Facility, supported by the Institutional Research Core Program and O’Neal Comprehensive Cancer Center (NIH grant P30 CA013148), with additional funding from NIH grant S10 OD024978. We are grateful to Dr. James Kizziah, Dr. Thomas Edwards, Dr. Tara Fox, Dr. Adam Wier, and Dr. Zhiqing Wang for assisting with the screening or data collection. We’d like to thank Dr. Ed Egelman, Dr. Mark Kreutzberger, and Dr. Daniel Bond for helpful discussions in the preparation of the manuscripts. The work in F.W. laboratory was supported by NIH grant GM138756, DoE grant SC0024303, and pilot grant from the UAB global center for craniofacial oral and dental disorders (GC-CODED). The work in the H.W. laboratory was supported by NIH grants R01 DE022350, R01 DE028329 and T90 DE030859.

## Author Contributions

J.L.F. and H.Z. performed sample preparation. J.L.F., H.A.P., S.K.H. and S.T.R. performed microscopy image analysis. H.Z. performed sequencing experiments to confirm flagellin sequences. N.F.B. and F.W. performed outer domains bioinformatic analysis. F.W. and H.W. obtained funding and supervised the research. F.W. wrote the manuscript with input from all authors.

## Data availability

The three-dimensional reconstructions have been deposited in the Electron Microscopy Data Bank with accession codes EMD-43829 (*C. gilardii* flagellum), EMD-43830 (*S. maltophilia* flagellum), and EMD-43868 (*G. thiophilus* flagellum). The atomic models have been deposited in the Protein Data Bank with accession codes 9ATB (*C. gilardii* flagellum), 9ATL (*S. maltophilia* flagellum), and 9AU6 (*G. thiophilus* flagellum).

## Supplementary Material

**Supplementary Figure 1.**
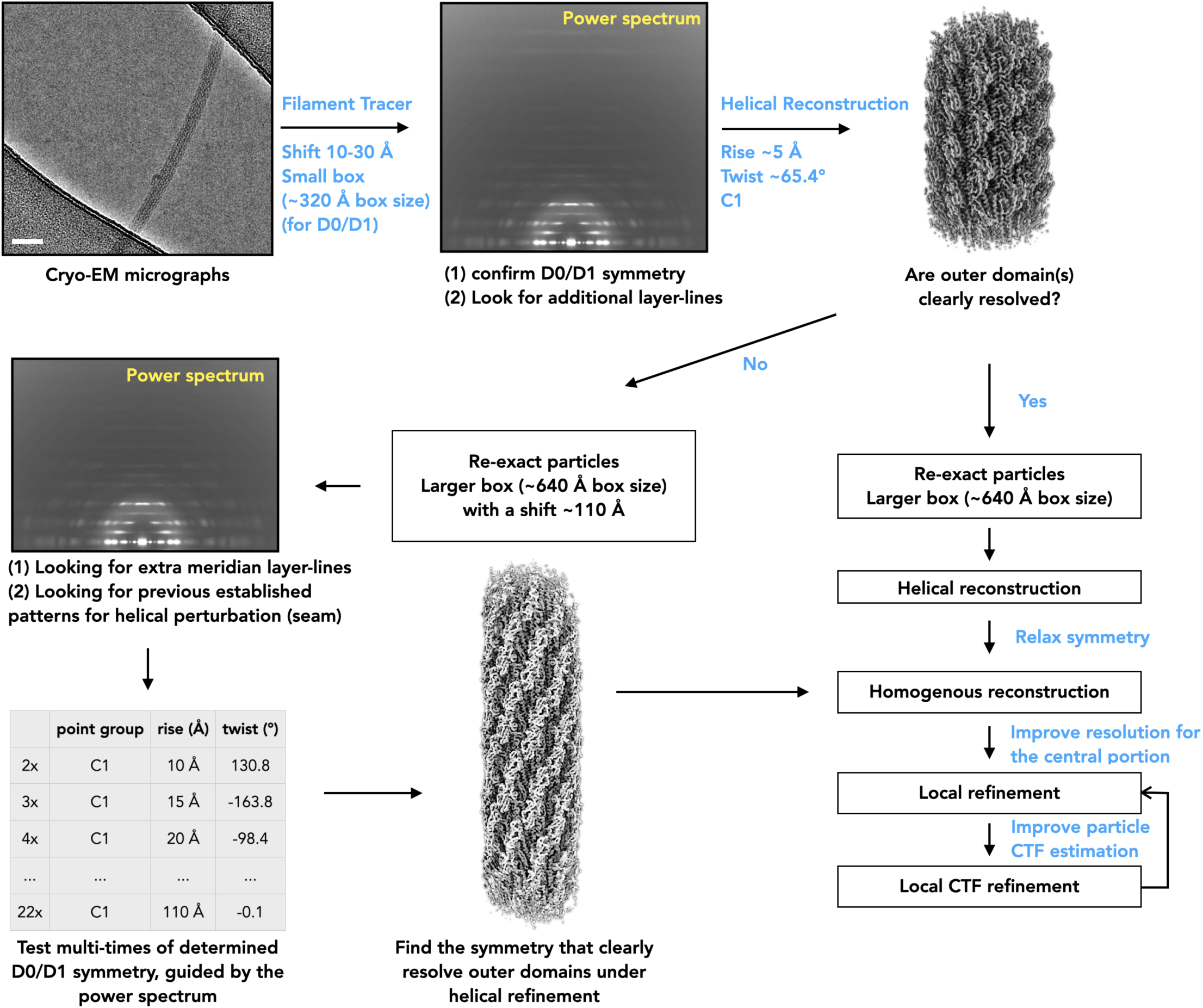
A general workflow for 3D reconstructions of supercoiled bacterial flagellar filaments. The micrographs in this study are captured at pixel size around 1 Å/px. The symmetry listed in the table are estimates and the number applied should be dependent on the symmetry refined in early helical refinement jobs.

**Supplementary Figure 2.**
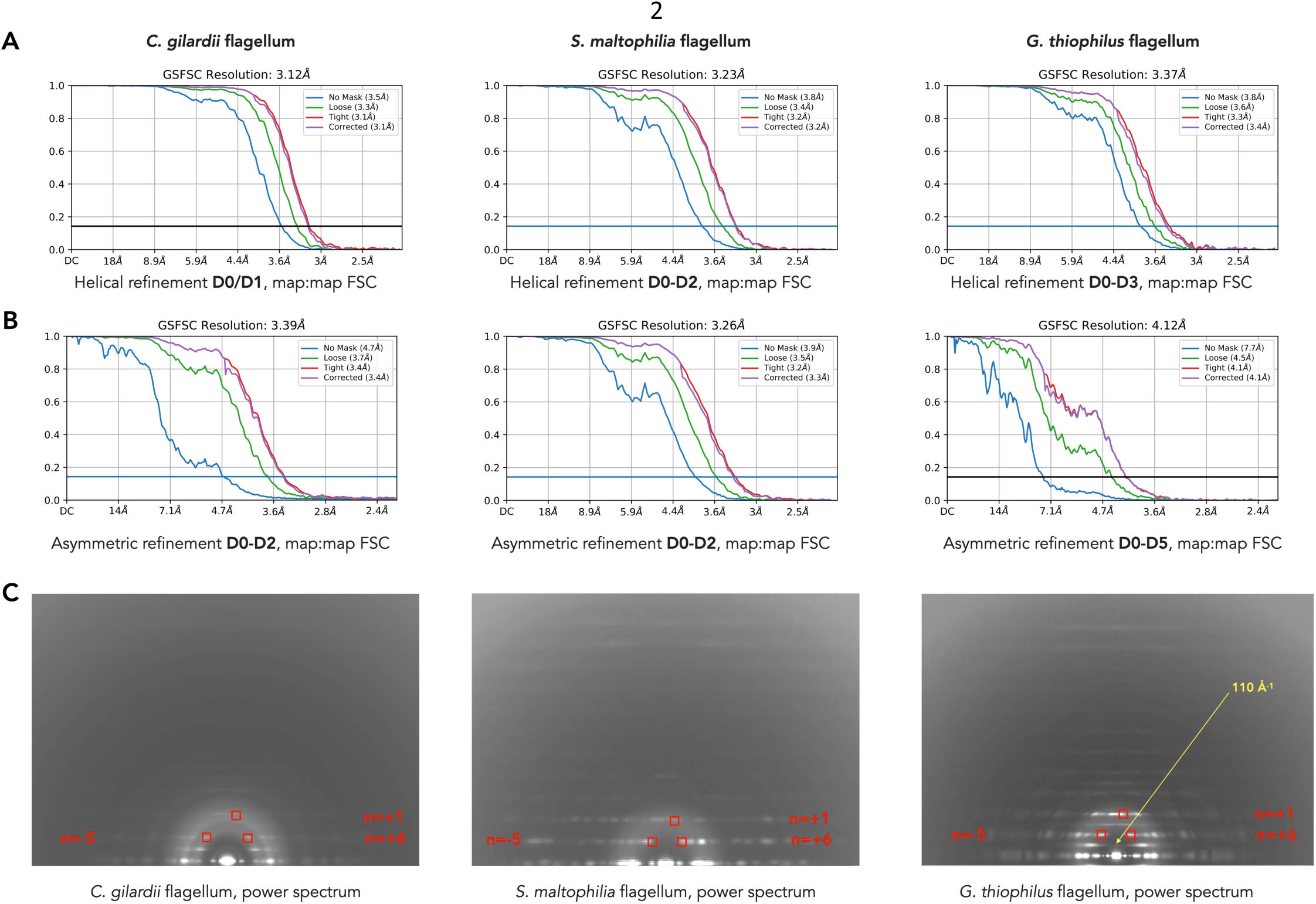
Fourier Shell Correlation (FSC) calculations and average power spectra of three flagellar filaments. (A) The map:map “gold-standard FSC” curves for initial 320-pixel box helical reconstruction in three flagellar filament datasets, applying the canonical D0/D1 symmetry (B) The map:map “gold-standard FSC” curves for the final local refinement using the 640-pixel box, relaxing the helical symmetry. (C) Average power spectra of particles behind 2D class average of *C. gilardii* flagellum (left), *S. maltophilia* flagellum (middle), and *G. thiophilus* flagellum (right). The Bessel orders of the layer lines from canonical flagellar D0/D1 domains are labeled in red.

**Supplementary Figure 3.**
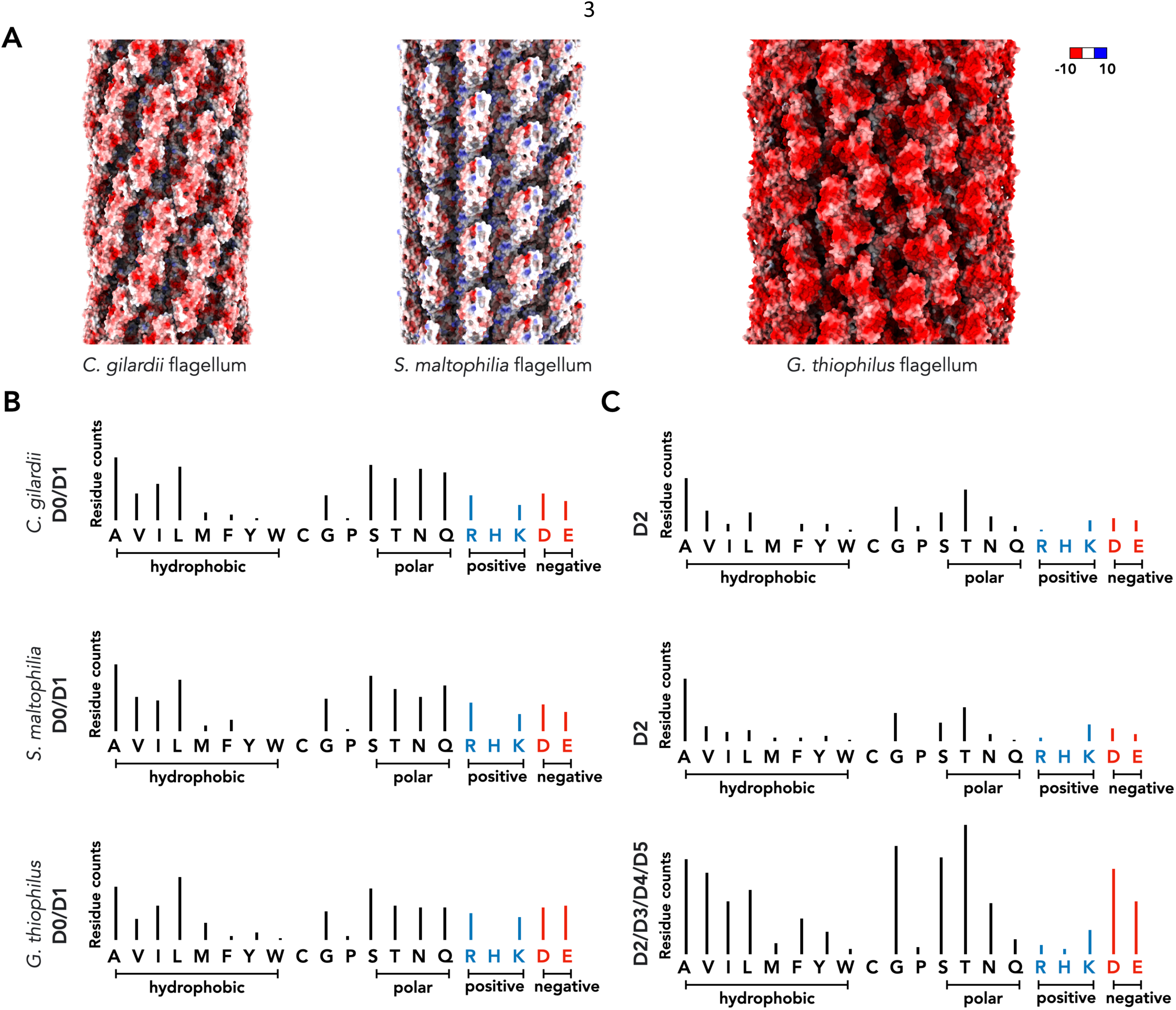
Electrostatic surfaces and amino acid components of three flagellar filaments. (A) The coulombic electrostatic potential calculations for *C. gilardii* flagellum (left), *S. maltophilia* flagellum (middle), and *G. thiophilus* flagellum (right). Scale bar displays the color-coding of the electrostatic potential in unit kcal·mol^-1^·*e*^-1^ at 298 K. (B) The amino acid components of D0/D1 domains in three flagellins. (C) The amino acid components of outer domains in three flagellins.

**Supplementary Figure 4.**
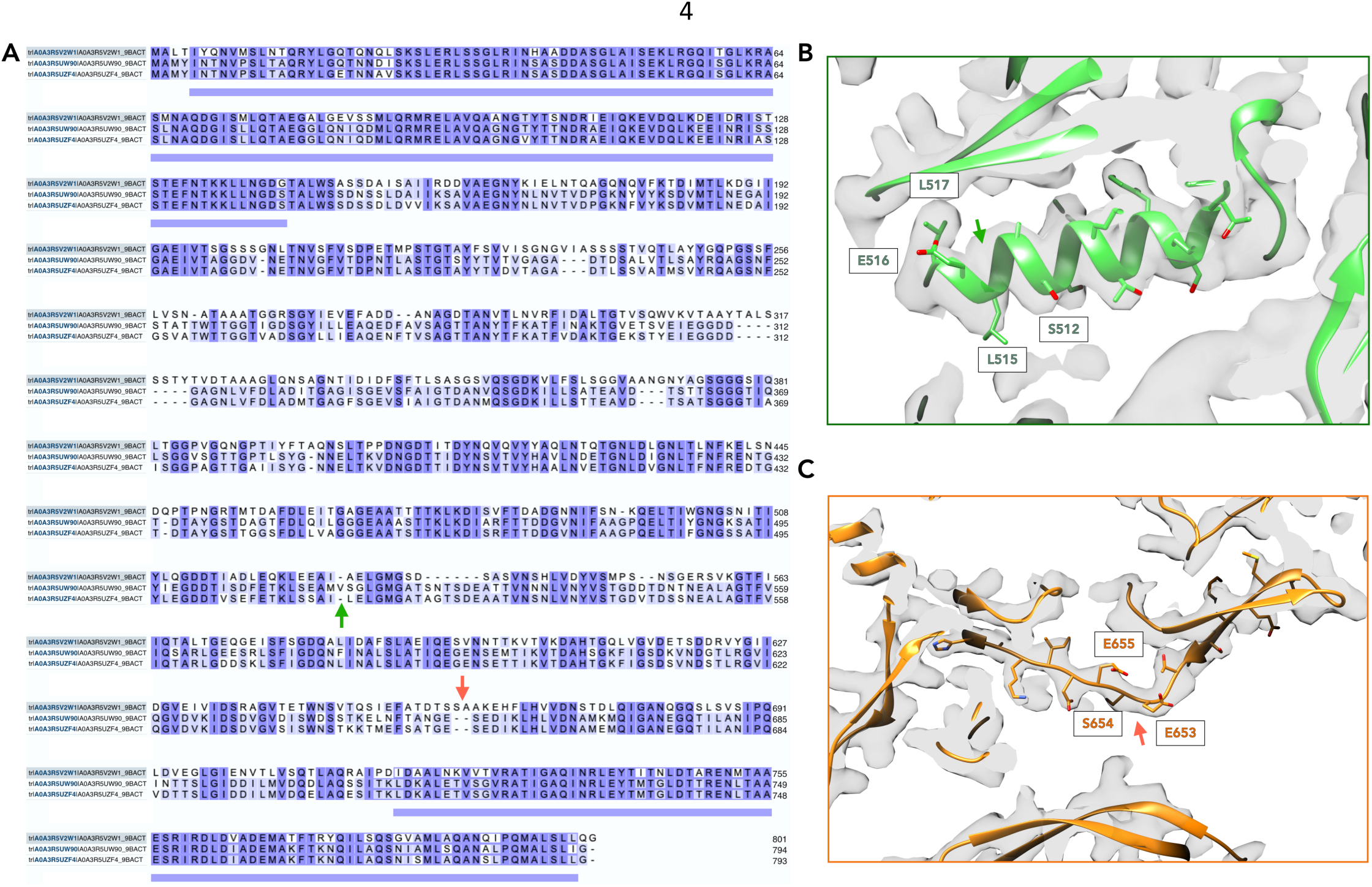
Identification of flagellin using cryo-EM map and structure modeling. **(A)** Sequence alignment of three similar flagellins from *G. thiophilus*. Their UniProt IDs are A0A3R5V2W1, A0A3R5UW90, and A0A3R5UZF4. A0A3R5UZF4 was identified to the correct flagellin, and example map areas for such identification is labeled using green and orange arrows. The corresponding map area was labeled using the same arrows in (**B**) and (**C**). **(B)** The map region where A0A3R5UW90 won’t be able to explain the cryo-EM density. If A0A3R5UW90 was the flagellin, it needs to insert one extra residue into the helix, and at L517, where clear site chain density exists, will be a glycine. This is clearly impossible. **(C)** The map region where A0A3R5V2W1 won’t be able to explain the cryo-EM map. If A0A3R5V2W1 was the flagellin, it needs to insert two residues into this well-resolved loop region, which is clearly impossible.

